# Improved Prediction of Smoking Status via Isoform-Aware RNA-seq Deep Learning Models

**DOI:** 10.1101/2020.09.09.290395

**Authors:** Zifeng Wang, Aria Masoomi, Zhonghui Xu, Adel Boueiz, Sool Lee, Tingting Zhao, Michael Cho, Edwin K. Silverman, Craig Hersh, Jennifer Dy, Peter J Castaldi

## Abstract

Most predictive models based on gene expression data do not leverage information related to gene splicing, despite the fact that splicing is a fundamental feature of eukaryotic gene expression. Cigarette smoking is an important environmental risk factor for many diseases, and it has profound effects on gene expression. Using smoking status as a prediction target, we developed deep neural network predictive models using gene, exon, and isoform level quantifications from RNA sequencing data in over 2,000 subjects in the COPDGene Study. We observed that models using exon and isoform quantifications clearly outperformed gene-level models when using data from 5 genes from a previously published five gene prediction model. Whereas the test set performance of the previously published model was 0.82 in the original publication, our exon-based models including an exon-to-isoform mapping layer achieved a test set AUC of 0.88 using data from the same 5 genes and an AUC of 0.94 using a larger set of exon quantifications. Isoform variability is an important source of latent information in RNA-seq data that can be used to improve clinical prediction models.

**Author summary:** Predictive models based on gene expression are already a part of medical decision making for selected situations such as early breast cancer treatment. Most of these models are based on measures that do not capture critical aspects of gene splicing, but with RNA sequencing it is possible to capture some of these aspects of alternative splicing and use them to improve clinical predictions. Building on previous models to predict cigarette smoking status, we show that measures of alternative splicing significantly improve the accuracy of these predictive models.

## Introduction

Smoking is the most important environmental risk factor for a wide range of diseases including cardiovascular disease, lung cancer, chronic obstructive pulmonary disease (COPD). Smoking increases the risk for these diseases through a variety of mechanisms including selective activation and repression of distinct aspects of the inflammatory response [1].

A meta-analysis of blood gene expression arrays from 5,376 current and former smokers identified 1,270 smoking-associated differentially expressed genes that were significantly enriched in immune-related processes including T-cell activation [2]. However, it is challenging to characterize the effects of cigarette smoking on splicing and differential isoform usage due to technical challenges in measuring isoform expression levels. Using RNA-seq combined with novel isoform reconstruction algorithms, we have shown that smoking causes widespread differential isoform and exon usage in addition to overall gene-level expression changes [3].

A five gene expression-based predictive model for smoking was previously proposed by Beineke et. al [4] with an AUC of 0.82, indicating that there is still room for improvement in predictive performance for expression-based prediction tools for current smoking status. Using blood RNA-seq data from 2,557 subjects in the COPDGene Study, we explored the relative utility of expression measures at the gene, exon, and isoform level using deep learning models tailored specifically to account for patterns of alternative splicing induced by smoking. We hypothesized that since smoking alters patterns of exon and isoform usage, greater predictive accuracy could be obtained by using exon and isoform-level quantifications to predict smoking status.

## Materials and methods

### Subject enrollment and data collection

This study includes 2,557 subjects from the COPDGene Study. COPDGene has been previously described [5]. Self-identified non-Hispanic whites (NHW) and African Americans (AA) between the ages of 45 and 80 years with a minimum of 10 pack-years lifetime smoking history were enrolled at 21 centers across the United States. COPDGene conducted two study visits approximately five years apart, and the ten-year visits are being completed. Starting at the second study visit, complete blood count (CBC) data and PaxGene RNA tubes were collected. Smoking history was ascertained by self-report. Participants defined as current smokers answered yes to the question “Do you smoke cigarettes now (as of one month ago?)”. Institutional review board approval and written informed consent were obtained.

### Total RNA extraction

Total RNA was extracted from PAXgene Blood RNA tubes using the Qiagen PreAnalytiX PAXgene Blood miRNA Kit (Qiagen, Valencia, CA). The extraction protocol was performed either manually or with the Qiagen QIAcube extraction robot according to the company’s standard operating procedure.

### cDNA library construction and sequencing

Globin reduction and cDNA library preparation for total RNA was performed with the Illumina TruSeq Stranded Total RNA with Ribo-Zero Globin kit (Illumina, Inc., San Diego, CA). Library quality control included quantification with picogreen, size analysis on an Agilent Bioanalyzer or Tapestation 2200 (Agilent, Santa Clara, CA), and qPCR quantitation against a standard curve. 75 bp paired end reads were generated on Illumina sequencers. Samples were sequenced to an average depth of 20 million reads. All sequenced samples had RIN ¿ 6.

### Sequencing read alignment, quality control and expression quantification

Reads were trimmed of TruSeq adapters using Skewer with default parameters [6]. Trimmed reads were aligned to the GRCh38 genome using the STAR aligner [7]. Quality control was performed using the FastQC and RNA-SeQC programs [8]. Samples were included for subsequent analysis if they had >10 million total reads, >80% of reads mapped to the reference genome, XIST and Y chromosome expression was consistent with reported gender, < 10% of R1 reads in the sense orientation, Pearson correlation ≥ 0.9 with samples in the same library construction batch, and concordant genotype calls between variants called from RNA sequencing reads and DNA genotyping.

Gene and transcript gene transfer file (GTF) annotation was downloaded from Biomart Ensembl database (Ensembl Genes release 94, GRCh38.p12 assembly) on October 21, 2018. We further derived exonic parts GTF annotation by breaking exons into disjoint parts sharing a common set of transcripts within a single gene. Sequencing read counts on gene and exonic part level genomic features were obtained from featureCounts function in Rsubread package (v1.32.2). Isoform level expression quantifications were derived using the Salmon program (v0.12.0) and the tximport package (v1.10.0). The gene, isoform, and exon count data used for this analysis are available in GEO [26, 27] (accession number XXXXXX).

### Filtering, normalization, differential expression and usage analysis

Genomic features (genes, isoforms or exonic parts) were filtered for both features that had very low and very high expression. The filter used to remove low expressed features was to remove features where the average counts per million (CPM) was < 0.2 or the feature was not expressed at a CPM ¿ 50 in at least 50 subjects. Extreme highly expressed features were defined as features attaining a CPM > 50,000 in at least one but fewer than 50 subjects. Differences in sequencing depth and RNA library composition between subjects were normalized using the TMM procedure from edgeR package (v3.24.3). Counts were transformed to log2 CPM values and quantile-normalized to further remove systematic noise. To avoid overfitting, we limited our set of genes to those contained within a set of 1,270 smoking-associated differentially expressed genes that had been identified in a previous study using samples that did not overlap with this study [2].

### Data usage and model validation

We analyzed blood RNA-seq data from 2,557 subjects in the COPDGene Study. We randomly split the data into training, validation, and testing sets containing 1637, 407, 513 subjects respectively. Model optimization and hyperparameter tuning was performed in the training data using 5-fold cross-validation. A small set of high-performing models were further evaluated in the validation dataset, and the testing data were used only for evaluation of the final set of models after all parameters and hyperparameters were fixed. The testing data was held by a separate analysis group using a different computer system to avoid any possibility of inadvertent use of test data in the model building process.

### Model Training

For all experiments, we train each model for 40 epochs with batch size 256 using Adam optimizer with learning rate 0.0003, *β*_1_ = 0.9, *β*_2_ = 0.999 and a dropout rate of 0.2. We set the weight of the L1 constraint used in the Feature Selection layer to be 0.0005. Unless otherwise specified, model layers were fully connected and ReLU nonlinear activation function were used. All the deep learning models were implemented in TensorFlow (v12.0.0) and Keras (v2.2.4.)

To identify high-performing model architectures, we adopted a layer-by-layer incremental search strategy. We first explored the optimal number of nodes for a single layer network by performing grid search. We evaluated from 2 to 512 nodes in the first layer, increasing by a factor of 2 at each step, i.e., 2, 4, 8, …, 512. The number of nodes at each layer was selected based on cross-validation performance, and then an additional layer was added using the same grid search strategy for node number with the constraint that each subsequent layer would have fewer nodes than the preceding layer. This process was repeated until no further gain in performance was achieved.

### Implementation of Isoform Map and Feature Selection Layers

To incorporate prior knowledge regarding the relationship of exons to transcript isoforms, we implemented an Isoform Map Layer (IML) which takes exon feature as input and outputs estimated isoform feature. This specially-designed layer is based on a standard fully-connected layer with weight **W**. This layer encodes known exon to isoform relationships in a binary relationship matrix **R** such that if exon *i* is contained within isoform *j,* we set **R**_*ij*_ = 1, otherwise **R**_*ij*_ = 0. This layer takes the relationship matrix to perform element-wise multiplication with the learnable weight matrix **W**. Thus, only canonical exon to isoform relationships can contribute to the final model. Exon to isoform relationships were obtained from the Ensemble v94 GTF file.

The Feature Selection Layer (FSL) associates every input feature with a non-negative learnable weight using an L1 constraint and outputs a reweighted feature vector of the same size as the input feature vector. The weights represent each feature’s importance with respect to smoking status prediction.

### Baseline models and model comparisons

To assess the effectiveness of our method, we compared our method against the current method proposed by Beineke et al., which is a logistic regression model using the following five genes: *CLDND1, LRRN3, GOPC, LEF1, MUC1.* We apply the Beineke model on our data by exploring logistic regression with exon, isoform and gene inputs considering only these five genes. For models evaluating the full set of genes, we trained elastic net models as a baseline for comparison with the weights for the L1 and L2 norms set at 0.0005. We obtain the optimal set of parameters of elastic net by conducting grid search and find out the best performance on the validation set. All statistical tests of model performance were analyzed using data from the test set and performed using R version 3.6. Direct comparisons between models were performed using the deLong test implemented in the pROC package.

## Results

RNA-seq data from 2,557 current and former smokers in the COPDGene Study were used to develop and test the predictive models. Data were randomly split into training, validation, and testing data. The use of data for model training, selection, and testing are described in Fig 1. The characteristics of the subjects in these datasets are shown in Table 1.

**Fig 1.**
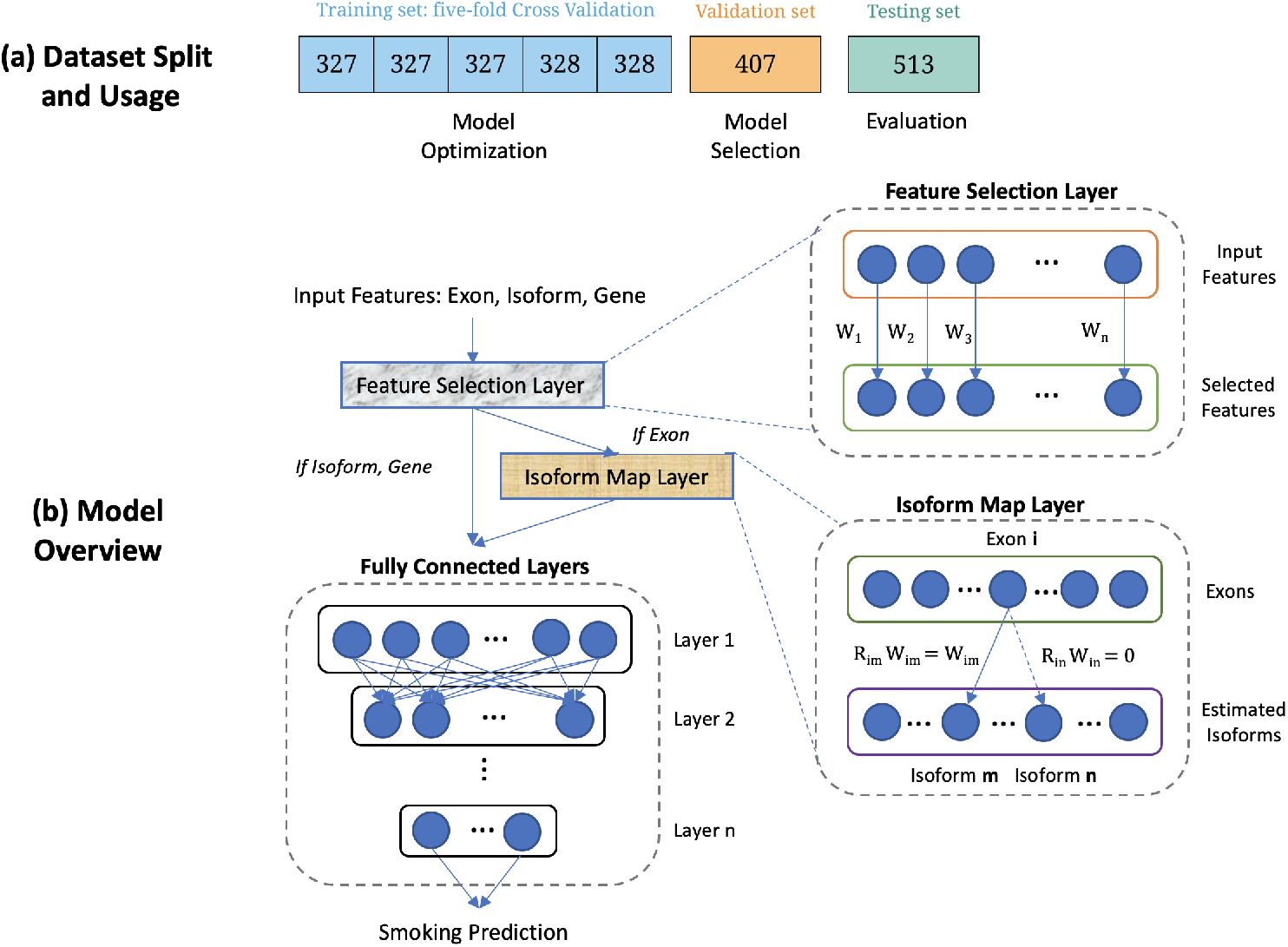
Visual Abstract. (a) Dataset split and usage. The number in each cell represents the number of subjects. The training set is equally split into 5 folds for deep learning model optimization (cross-validation for tuning the hyperparameters and architecture search in a deep learning model). The validation set is used to select the optimal model and the testing set is held out for performance evaluation. (b) Model overview. Our model consists of a Feature Selection Layer (FSL), an Isoform Map Layer (IML) (if the input feature is exon) and standard fully connected layers. FSL associates each input feature with a non-negative learnable weight, which represents the importance of features with respect to smoking status. IML encodes exon to isoform relationships via a binary matrix **R**, such that if exon *i* is contained within isoform *j*, we set **R**_*ij*_ = 1, otherwise **R**_*ij*_ = 0. By multiplying **R**_*ij*_ with corresponding learnable weights **W**, we only consider canonical exon to isoform relationships.

**Table 1.**
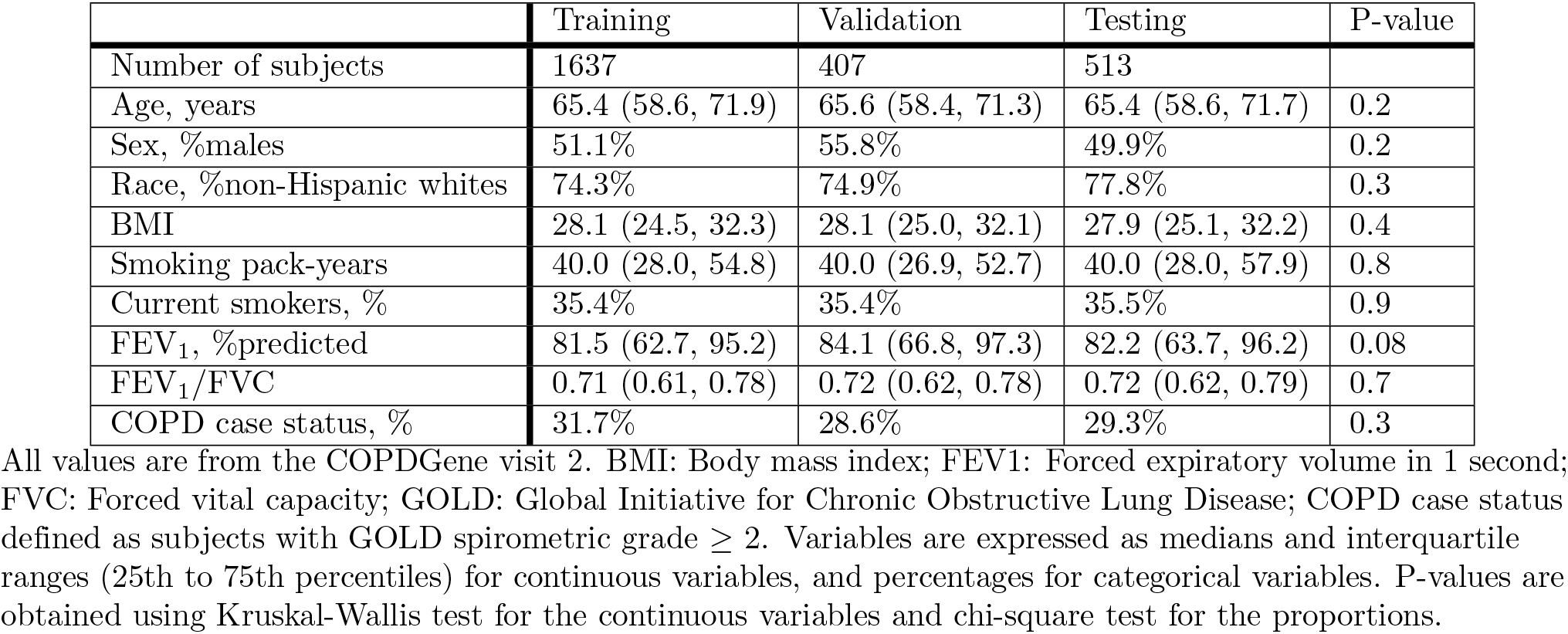
Characteristics of subjects.

### Validation and Further Development of the Beineke Model using Exon and Isoform Level Data

A microarray and RT-PCR based five gene expression model for smoking has been previously developed and shown to have a test set AUC of 0.82 [4]. To externally validate this model and establish a performance benchmark in our dataset, we constructed an initial set of models using gene, exon, and isoform expression from this set of genes. One of the genes, *MUC1* was expressed in our data at levels below our filtering threshold. We confirmed that this gene is also expressed in very low levels in whole blood RNA samples from the Genotype Tissue Expression Project, and subsequently based our models on the other four gene expression values. A logistic regression model using these four genes had an AUC of 0.76 and 0.78 in our validation and testing data (Table 2). We then trained two additional logistic regression models using exon counts and Salmon estimated isoform quantifications from these genes. As shown in Fig 2, the prediction performance in both validation and testing datasets was improved using both isoform (p=0.002) and exon level (p<0.001) quantifications, and exon data outperformed Salmon estimated isoform data (p=0.002). Notably, the best performing models used exon level data combined with an (exon-to-)Isoform Map Layer based on curated isoform data (i.e. Ensembl GTF) and a Feature Selection Layer, as described later.

**Fig 2.**
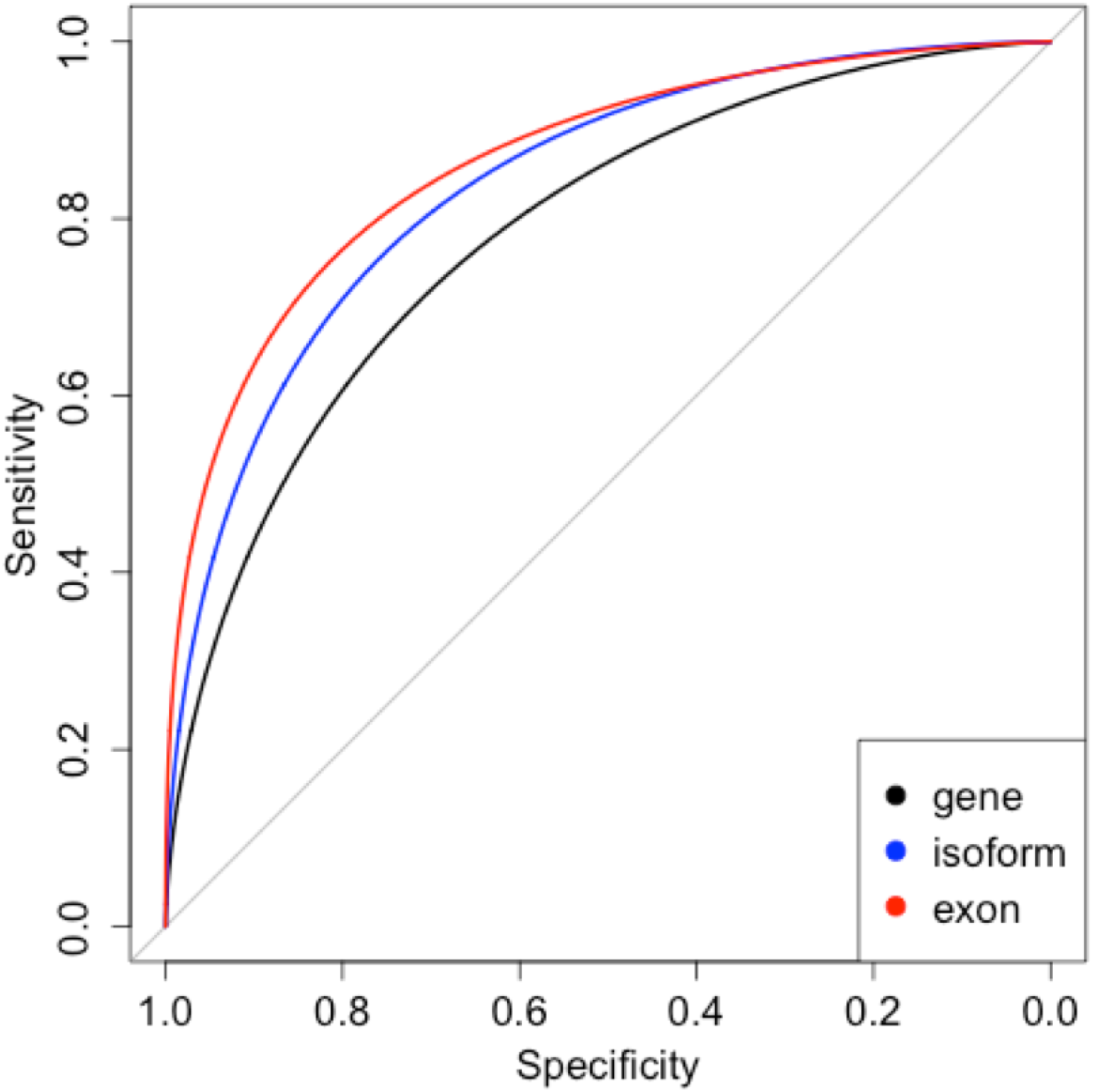
ROC curves in test data for the 4-gene modified Beineke model using gene (black), isoform (blue), and exon-level (red) quantifications. Isoform and exon-level data outperform gene-level data (Delong p=0.002 and <0.001, respectively).

**Fig 3.**
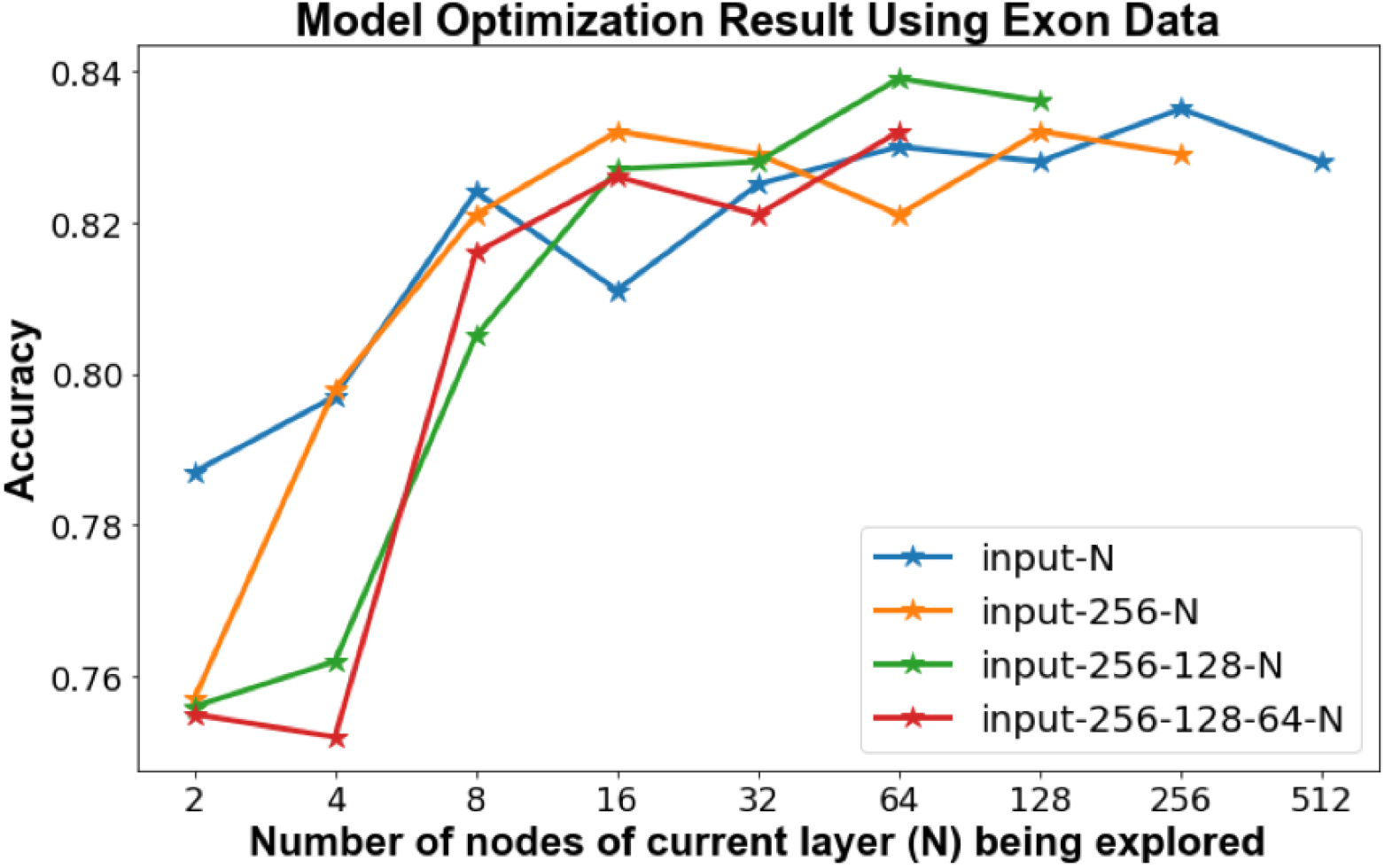
Cross-validation accuracy calculated during model optimization for exon-level data.

**Table 2.** Predictive performance of modified Beineke models using gene, isoform and exon-level expression data.

|  | Val - Accuracy | Val - AUC | Test - Accuracy | Test - AUC |
| --- | --- | --- | --- | --- |
| Gene | 0.698 | 0.758 | 0.743 | 0.780 |
| Isoform | 0.757 | 0.828 | 0.774 | 0.828 |
| Exon | 0.801 | 0.859 | 0.808 | 0.869 |
| Exon, IML-GTF | **0.828** | 0.876 | 0.825 | 0.870 |
| Exon, IML-GTF, FSL | **0.828** | **0.889** | **0.838** | **0.875** |

Val: validation data. AUC: area under the curve. IML-GTF: Isoform Map Layer containing information from Ensembl GTF file. FSL: Feature Selection Layer. Best results are shown in bold.

### Model Optimization Using A Larger Feature Set

Having obtained improved prediction performance using exon and isoform data from four genes in the Beineke model, we then constructed models using a much larger set of features. Of the 1,270 genes that were significantly associated with current smoking from the meta-analysis by Huan et al. [2], 1,079 were expressed at levels high enough to be analyzed in our RNA-seq data. These genes contained 6,196 isoforms and 19,027 exons present in our data, and we constructed separate deep learning models using gene, isoform, and exon level data. As expected, the best models for isoform and exon data had a larger number of nodes (256-128-64) than the gene level model (128-64-32). Maximal accuracy was observed with three layers, and the best performance was achieved with exon level quantifications (Fig 4).

**Fig 4.**
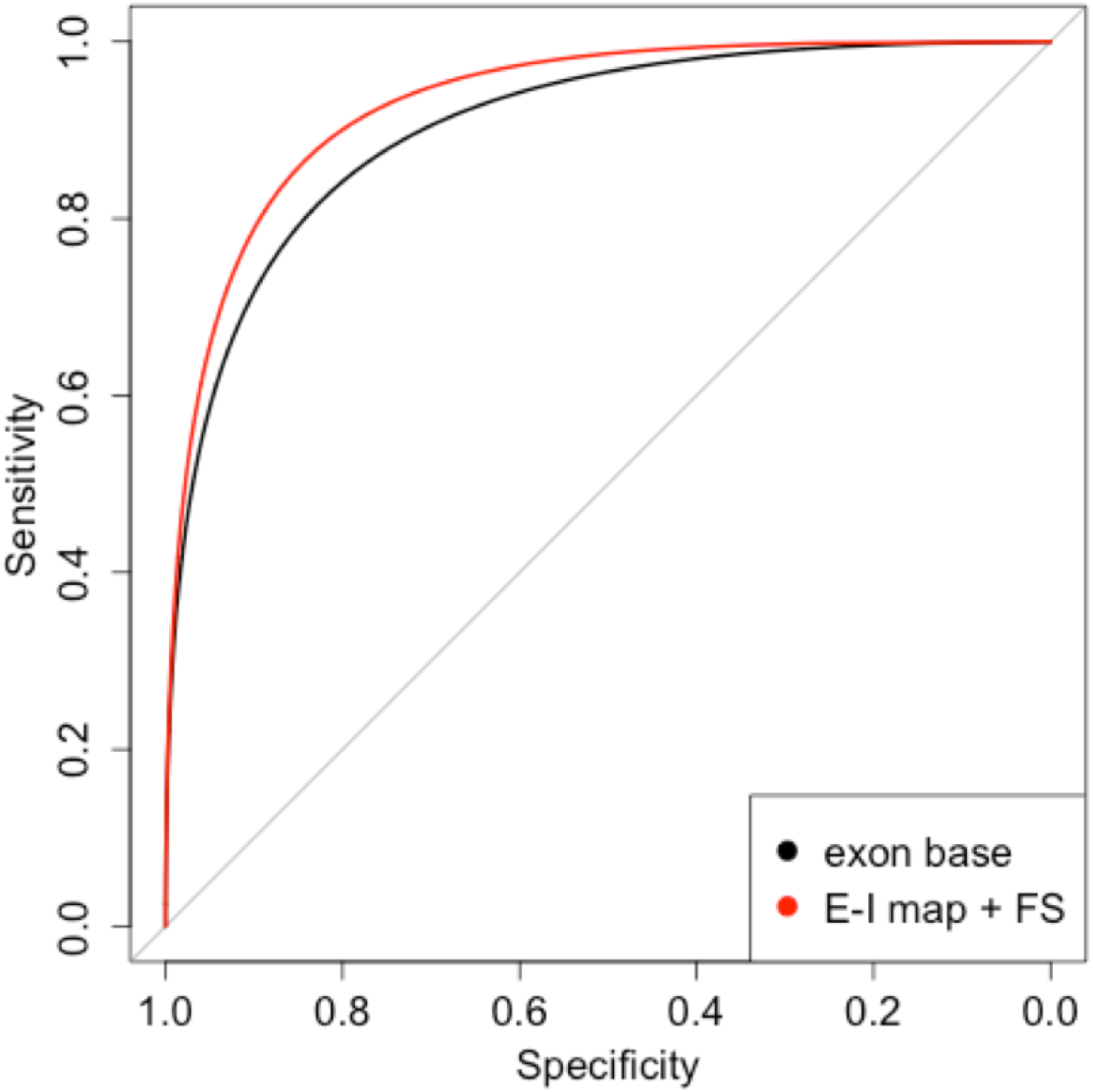
ROC curves in test data for the deep learning base exon model (black) and the model including the exon-isoform mapping and feature selection layers (red) which has significantly better performance (Delong test p=0.02).

### Improved Prediction through Exon-to-Isoform Mapping and Feature Selection Layers

We hypothesized that the performance of exon-based prediction models would be improved by incorporating relationships between exons and isoforms. Using known exon to isoform relationships from the Ensembl version 94 GTF file, we introduced a deep learning layer (IML) that encoded these connections between exons and isoforms (Figure 1), and we observed improved predictive performance in cross-validation and in test data (Table 3). For comparison, we also compared these models to models that incorporated a fully connected layer between exons and isoforms, but this model was far more complex and failed to converge.

We then explored whether the addition of an integrated feature selection layer (FSL) would further improve performance by introducing an additional layer that assigns a non-negative weight for each input feature, and we observed an incremental increase in performance. When we compared the performance of this model to the base exon model, the performance was significantly improved (p=0.02 in test data, Figure 4).

**Table 3.** Predictive performance of various models using exon-level data, including elastic net for comparison.

|  | Val - Accuracy | Val - AUC | Test - Accuracy | Test - AUC |
| --- | --- | --- | --- | --- |
| Exon, Elastic Net | 0.821 | 0.861 | 0.774 | 0.903 |
| Exon Base | 0.813 | 0.886 | 0.842 | 0.913 |
| Exon, IML-GTF | 0.843 | 0.905 | 0.854 | 0.924 |
| Exon, IML-GTF, FSL | **0.860** | **0.916** | **0.869** | **0.935** |

Val: validation data. AUC: area under the curve. IML-GTF: Isoform Map Layer containing information from GTF file. FSL: Feature Selection Layer. Best results are shown in bold.

## Discussion

Deep learning models applied to blood RNA-seq data provide more accurate prediction of current smoking status than previously published models. In testing data, our models achieved an AUC >0.9 compared to a replication AUC of 0.81 for the previously established 5-gene model. Much of this improvement is due to the use of exon rather than gene expression levels coupled with the use of a neural net layer encoding exon to isoform relationships. These findings improve our ability to identify environmental exposures from RNA-seq data, and they suggest that latent isoform information in RNA-seq data can be used to improve clinical predictions.

This paper describes for the first time how exon and isoform-level data from RNA-seq improve the accuracy of clinical prediction models, demonstrating a general approach by which gene expression predictive models may be improved. Eukaryotic genomes are characterized by complex gene structure and extensive alternative splicing that greatly expands the protein repertoire. Over 90% of human genes have multiple transcribed isoforms [9], isoform variability is clearly observable across tissues within the same individual [10], and isoform variability is an important contributor to human diseases [11, 12]. Focusing first on the previously published Beineke gene expression model, we demonstrate a notable increase in performance by substituting exon or isoform quantifications for the same set of genes used in the original model (AUC increase from 0.76 to 0.86). The best performance was achieved with exon data, not estimated isoform quantifications, which is likely due to inaccuracy in the estimation of full length isoforms from short-read RNA-seq.

We were able to further improve our model by encoding known exon-isoform relationships in one of the layers of the neural network, which we refer to as the isoform mapping layer. This is in line with other applications of machine learning to biological data that have found improved performance for algorithms that can incorporate prior biological knowledge, such as the use of known gene-interaction networks to improve the performance of clustering methods [13,14]. Since our current catalog of human isoform variability is incomplete, as this knowledge increases the value of isoform mapping layers that depend on prior knowledge will also increase. In addition, with the growing use of long read sequencing, these highly accurate isoform quantifications can be used directly as inputs to predictive models. Our data suggest that this will lead to further improvements in predictive accuracy for models based on RNA expression.

Gene expression prediction models for current smoking status are useful for multiple reasons. First, existing smoking biomarkers have good but not ideal predictive performance. In clinical practice, determination of smoking status is primarily done by patient self-report, and in instances where biochemical validation is necessary this is done via measures of nicotine metabolites, such as cotinine, in blood, urine, or saliva. While it may seem straightforward to determine smoking status, in practice it is difficult to ascertain smoking status with complete certainty for multiple reasons. Individuals may not accurately report their smoking behavior, and biochemical tests can yield false positives when individuals are exposed to nicotine in the absence of cigarette use, as can occur with the use of nicotine replacement therapy or electronic nicotine delivery devices (e-cigarettes). A systematic review of the performance of various cotinine cutoffs with respect to self-report of smoking status reported performance in the range of 70-90% sensitivity with specificity levels of 98% [15]. While our models outperformed previous models based on expression data, they did not perform as well as cotinine with respect to predicting self-reported smoking status. Thus, from the standpoint of clinical biomarkers for smoking status, nicotine metabolites such as cotinine remain the gold standard. Our model is best used for situations where gene expression data are available, but cotinine measures are not.

Another important application for transcriptome-based predictive models is to infer smoking status when only gene expression data are available. This is important because smoking has a strong effect on gene expression and therefore can be a confounder of gene expression studies, particularly in situations where smoking is confounded with specific disease states. In this scenario, use of a previously defined model for to infer smoking status may allow for more accurate detection of disease-related gene expression signals, even when the smoking status of subjects has not been directly measured.

The strengths of this study are the large sample size of subjects with blood RNA-seq data, and the ability to assess our predictive models in two sets of independent test data. We assessed deep learning based method which has provided superior predictive performance in multiple contexts, and we assessed the predictive utility of novel aspects of RNA expression which have not been extensively studied in the prediction context. Limitations of this study are that smoking status was determined by self-report only, and cotinine measures were not available.

## Conclusion

In summary, the use of exon-level quantifications in combination with an exon-to-isoform mapping layer produced predictive models with superior ability to predict current smoking status relative to previously published models from gene expression data. While these models still do not outperform gold-standard metabolite biomarkers of smoking, they can be of use in studies where such biomarkers are not available. Finally, these findings are proof-of-concept that incorporating isoform-level information into predictive models improves the ability to predict clinical outcomes. As the quality of isoform quantification improves from isoform inference algorithms and long-read sequencing, it is reasonable to expect that the performance of RNA-based predictive models will also improve.

## Supporting information

## Acknowledgments

The project described was supported by Award Numbers U01 HL089897, U01 HL089856, R01 HL124233, and R01 HL147326 from the National Heart, Lung, and Blood Institute and the FDA Center for Tobacco Products (CTP). The content is solely the responsibility of the authors and does not necessarily represent the official views of the NIH or the Food and Drug Administration.

COPD Foundation Funding The COPDGene® project is also supported by the COPD Foundation through contributions made to an Industry Advisory Board comprised of AstraZeneca, Boehringer-Ingelheim, Genentech, GlaxoSmithKline, Novartis, and Sunovion.

## COPDGene® Investigators – Core Units

Administrative Center: James D. Crapo, MD (PI); Edwin K. Silverman, MD, PhD (PI); Barry J. Make, MD; Elizabeth A. Regan, MD, PhD

Genetic Analysis Center: Terri Beaty, PhD; Ferdouse Begum, PhD; Peter J. Castaldi, MD, MSc; Michael Cho, MD; Dawn L. DeMeo, MD, MPH; Adel R. Boueiz, MD; Marilyn G. Foreman, MD, MS; Eitan Halper-Stromberg; Lystra P. Hayden, MD, MMSc; Craig P. Hersh, MD, MPH; Jacqueline Hetmanski, MS, MPH; Brian D. Hobbs, MD; John E. Hokanson, MPH, PhD; Nan Laird, PhD; Christoph Lange, PhD; Sharon M. Lutz, PhD; Merry-Lynn McDonald, PhD; Margaret M. Parker, PhD; Dmitry Prokopenko, Ph.D; Dandi Qiao, PhD; Elizabeth A. Regan, MD, PhD; Phuwanat Sakornsakolpat, MD; Edwin K. Silverman, MD, PhD; Emily S. Wan, MD; Sungho Won, PhD

Imaging Center: Juan Pablo Centeno; Jean-Paul Charbonnier, PhD; Harvey O. Coxson, PhD; Craig J. Galban, PhD; MeiLan K. Han, MD, MS; Eric A. Hoffman, Stephen Humphries, PhD; Francine L. Jacobson, MD, MPH; Philip F. Judy, PhD; Ella A. Kazerooni, MD; Alex Kluiber; David A. Lynch, MB; Pietro Nardelli, PhD; John D. Newell, Jr., MD; Aleena Notary; Andrea Oh, MD; Elizabeth A. Regan, MD, PhD;

James C. Ross, PhD; Raul San Jose Estepar, PhD; Joyce Schroeder, MD; Jered Sieren; Berend C. Stoel, PhD; Juerg Tschirren, PhD; Edwin Van Beek, MD, PhD; Bram van Ginneken, PhD; Eva van Rikxoort, PhD; Gonzalo Vegas Sanchez-Ferrero, PhD; Lucas Veitel; George R. Washko, MD; Carla G. Wilson, MS;

PFT QA Center, Salt Lake City, UT: Robert Jensen, PhD

Data Coordinating Center and Biostatistics, National Jewish Health, Denver, CO: Douglas Everett, PhD; Jim Crooks, PhD; Katherine Pratte, PhD; Matt Strand, PhD; Carla G. Wilson, MS

Epidemiology Core, University of Colorado Anschutz Medical Campus, Aurora, CO: John E. Hokanson, MPH, PhD; Gregory Kinney, MPH, PhD; Sharon M. Lutz, PhD; Kendra A. Young, PhD

Mortality Adjudication Core: Surya P. Bhatt, MD; Jessica Bon, MD; Alejandro A. Diaz, MD, MPH; MeiLan K. Han, MD, MS; Barry Make, MD; Susan Murray, ScD; Elizabeth Regan, MD; Xavier Soler, MD; Carla G. Wilson, MS

Biomarker Core: Russell P. Bowler, MD, PhD; Katerina Kechris, PhD; Farnoush Banaei-Kashani, Ph.D

COPDGene® Investigators – Clinical Centers

Ann Arbor VA: Jeffrey L. Curtis, MD; Perry G. Pernicano, MD

Baylor College of Medicine, Houston, TX: Nicola Hanania, MD, MS; Mustafa Atik, MD; Aladin Boriek, PhD; Kalpatha Guntupalli, MD; Elizabeth Guy, MD; Amit Parulekar, MD;

Brigham and Women’s Hospital, Boston, MA: Dawn L. DeMeo, MD, MPH; Alejandro A. Diaz, MD, MPH; Lystra P. Hayden, MD; Brian D. Hobbs, MD; Craig Hersh, MD, MPH; Francine L. Jacobson, MD, MPH; George Washko, MD

Columbia University, New York, NY: R. Graham Barr, MD, DrPH; John Austin, MD; Belinda D’Souza, MD; Byron Thomashow, MD

Duke University Medical Center, Durham, NC: Neil MacIntyre, Jr., MD; H. Page McAdams, MD; Lacey Washington, MD

Grady Memorial Hospital, Atlanta, GA: Eric Flenaugh, MD; Silanth Terpenning, MD

HealthPartners Research Institute, Minneapolis, MN: Charlene McEvoy, MD, MPH; Joseph Tashjian, MD

Johns Hopkins University, Baltimore, MD: Robert Wise, MD; Robert Brown, MD; Nadia N. Hansel, MD, MPH; Karen Horton, MD; Allison Lambert, MD, MHS; Nirupama Putcha, MD, MHS

Lundquist Institute for Biomedical Innovationat Harbor UCLA Medical Center, Torrance, CA: Richard Casaburi, PhD, MD; Alessandra Adami, PhD; Matthew Budoff, MD; Hans Fischer, MD; Janos Porszasz, MD, PhD; Harry Rossiter, PhD; William Stringer, MD

Michael E. DeBakey VAMC, Houston, TX: Amir Sharafkhaneh, MD, PhD; Charlie Lan, DO

Minneapolis VA: Christine Wendt, MD; Brian Bell, MD; Ken M. Kunisaki, MD, MS

National Jewish Health, Denver, CO: Russell Bowler, MD, PhD; David A. Lynch, MB

Reliant Medical Group, Worcester, MA: Richard Rosiello, MD; David Pace, MD

Temple University, Philadelphia, PA: Gerard Criner, MD; David Ciccolella, MD; Francis Cordova, MD; Chandra Dass, MD; Gilbert D’Alonzo, DO; Parag Desai, MD; Michael Jacobs, PharmD; Steven Kelsen, MD, PhD; Victor Kim, MD; A. James Mamary, MD; Nathaniel Marchetti, DO; Aditi Satti, MD; Kartik Shenoy, MD; Robert M. Steiner, MD; Alex Swift, MD; Irene Swift, MD; Maria Elena Vega-Sanchez, MD

University of Alabama, Birmingham, AL: Mark Dransfield, MD; William Bailey, MD; Surya P. Bhatt, MD; Anand Iyer, MD; Hrudaya Nath, MD; J. Michael Wells, MD

University of California, San Diego, CA: Douglas Conrad, MD; Xavier Soler, MD, PhD; Andrew Yen, MD

University of Iowa, Iowa City, IA: Alejandro P. Comellas, MD; Karin F. Hoth, PhD; John Newell, Jr., MD; Brad Thompson, MD

University of Michigan, Ann Arbor, MI: MeiLan K. Han, MD MS; Ella Kazerooni, MD MS; Wassim Labaki, MD MS; Craig Galban, PhD; Dharshan Vummidi, MD

University of Minnesota, Minneapolis, MN: Joanne Billings, MD; Abbie Begnaud, MD; Tadashi Allen, MD

University of Pittsburgh, Pittsburgh, PA: Frank Sciurba, MD; Jessica Bon, MD; Divay Chandra, MD, MSc; Carl Fuhrman, MD; Joel Weissfeld, MD, MPH

University of Texas Health, San Antonio, San Antonio, TX: Antonio Anzueto, MD; Sandra Adams, MD; Diego Maselli-Caceres, MD; Mario E. Ruiz, MD; Harjinder Singh

## References

1. Arnson Y, Shoenfeld Y, Amital H. Effects of tobacco smoke on immunity, inflammation and autoimmunity. 2010;34(3):J258–65.

2. Huan T, Joehanes R, Schurmann C, Schramm K, Pilling LC, Peters MJ, et al. A whole-blood transcriptome meta-analysis identifies gene expression signatures of cigarette smoking. Human molecular genetics. 2016;25(21):4611–4623.

3. Parker MM, Chase RP, Lamb A, Reyes A, Saferali A, Yun JH, et al. RNA sequencing identifies novel non-coding RNA and exon-specific effects associated with cigarette smoking. BMC medical genomics. 2017;10(1):58.

4. Beineke P, Fitch K, Tao H, Elashoff MR, Rosenberg S, Kraus WE, et al. A whole blood gene expression-based signature for smoking status. BMC medical genomics. 2012;5(1):58.

5. Regan EA, Hokanson JE, Murphy JR, Make B, Lynch DA, Beaty TH, et al. Genetic epidemiology of COPD (COPDGene) study design. COPD: Journal of Chronic Obstructive Pulmonary Disease. 2010;7(1):32–43.

6. Jiang H, Lei R, Ding SW, Zhu S. Skewer: a fast and accurate adapter trimmer for next-generation sequencing paired-end reads. BMC Bioinformatics. 2014;15(1):182.

7. Dobin A, Dobin A, Davis CA, Schlesinger F, Schlesinger F, Drenkow J, et al. STAR: ultrafast universal RNA-seq aligner. Bioinformatics. 2013;29(1):15–21.

8. DeLuca DS, Levin JZ, Sivachenko A, Fennell T, Nazaire MD, Williams C, et al. RNA-SeQC: RNA-seq metrics for quality control and process optimization. Bioinformatics. 2012;28(11):1530–1532.

9. Wang ET, Sandberg R, Luo S, Khrebtukova I, Zhang L, Mayr C, et al. Alternative isoform regulation in human tissue transcriptomes. Nature. 2008;456(7221):470–476.

10. Reyes A, Huber W. Alternative start and termination sites of transcription drive most transcript isoform differences across human tissues. Nucleic Acids Research. 2018;46(2):582–592.

11. Scotti MM, Swanson MS. RNA mis-splicing in disease. Nature Reviews Genetics. 2016;17(1):19–32.

12. Li YI, van de Geijn B, Raj A, Knowles DA, Petti AA, Golan D, et al. RNA splicing is a primary link between genetic variation and disease. Science. 2016;352(6285):600–604.

13. Chang Y, Glass K, Liu YY, Silverman E, Crapo JD, Tal-Singer R, et al. COPD subtypes identified by network-based clustering of blood gene expression. Genomics. 2016;107(2-3):51–58.

14. Hofree M, Ideker TG, Shen JP, Carter H, Gross A. Network-based stratification of tumor mutations. Nature Methods. 2013;10(11):1108–1115.

15. Kim S. Overview of Cotinine Cutoff Values for Smoking Status Classification. International Journal of Environmental Research and Public Health. 2016;13(12):1236.

